# Severely lipoatrophic mice are hypermetabolic and hyperthermic under thermoneutral conditions in part due to an enhanced liver *de novo* lipogenesis

**DOI:** 10.64898/2026.06.18.733153

**Authors:** Álbert S. Peixoto, Caroline A. Lino, Bianca F. Leonardi, Érique Castro, Thayna S. Vieira, Julia V. França, Ana B. Pires, Natália M. Pessoa, Erika V. M. Pessoa, Marina A. Abe-Honda, Luciano P. Silva Júnior, Ana Carolina P. Baptista, Loreana Silveira, Maria Luiza E. Michalani, Mariana Mesquita, Samuel Santana, Erick M. Silveira, Luara B. Novaes, Adriano B. Chaves-Filho, Rafael J. Moreira, Tiago E. Oliveira, Helayne S. de Freitas, Camila N. Bezerra, William T. Festuccia

**Affiliations:** Institute of Biomedical Sciences, University of Sao Paulo, Sao Paulo, Brazil

**Keywords:** lipoatrophy/lipodystrophy, energy expenditure, *de novo* lipogenesis, thermoneutrality

## Abstract

White, beige and brown adipocytes store energy as lipids, secrete hormones and produce heat, playing an important role in the regulation of energy balance through not completely defined mechanisms. We investigate herein the impact of the almost complete absence of mature adipocytes (severe lipoatrophy) in the determination of energy balance (energy intake and expenditure) and homeothermy in mice. For this, mice with severe lipoatrophy induced by adipocyte deletion of peroxisome proliferator-activated receptor γ (PPARγ) (PPARγ flox adiponectin-Cre) and littermate controls (PPARγ flox) were evaluated for energy balance, thermoneutral zone, core body temperature, locomotor activity, and gene expression profiles at different ambient temperatures. Severely lipoatrophic mice are heavier, hypermetabolic and hyperphagic and feature a widened thermoneutral zone, lower ambulatory activity, and metabolic inflexibility at both 23 and 17°C, along with unstable thermal behavior characterized by hyperthermia at 30°C, normothermia at 23°C, and bouts of hypothermia at 17°C. Noteworthy, lipoatrophic mice hypermetabolism at 30°C is not due to thyroid hormones, impaired insulation or increased body and lean masses and is not altered by pharmacological blockade of either β-adrenergic receptor signaling with propranolol or skeletal muscle sarcoplasmic/endoplasmic reticulum Ca^2+^-ATPases (SERCA) and sarcolipin (SLN)-mediated calcium cycling with dantrolene, but is partially attenuated by pharmacological inhibition of acetyl-CoA carboxylase (ACC) and *de novo* lipogenesis with ND-630. In conclusion, severe lipoatrophy causes hypermetabolism and hyperthermia at 30°C partly through the activation of liver *de novo* fatty acid synthesis.

## 1. Introduction

White and brown adipocytes play major, but distinct roles in the determination of energy balance and metabolism. In addition to their endocrine function, white adipocytes mainly store/release lipids (energy) in periods of surplus and demand, respectively, while brown adipocytes generate heat through the uncoupling protein 1 (UCP-1)-mediated dissipation of mitochondrial proton gradient, but also the activation of ATP consuming-futile metabolic cycles. Supporting such important adipocyte role in the regulation of metabolism, a severe reduction in the number of white and brown adipocytes, and thus, white and brown adipose tissues (WAT and BAT, respectively) masses, collectively known as lipodystrophy/lipoatrophy, is associated with numerous metabolic complications including hyperglycemia, hyperinsulinemia, insulin resistance, hyperlipidemia, organomegaly, among others [1–3].

Conversely to these phenotypes, the impact of lipoatrophy on energy expenditure in both humans and rodents has yet to be fully elucidated with conflicting reports showing either upregulation, or reduction or no alteration. In humans, for instance, several studies reported a marked increase in resting energy expenditure in lipoatrophic patients either living with human immunodeficiency virus (HIV) (reviewed in [4]) or bearing genetic forms of lipodystrophy, such phenotype that persisted even after correction by lean mass [5]. Nevertheless, other reports found no changes in energy expenditure in lipoatrophic patients either before or after correction by lean mass [6,7].

A similar conflicting scenario is found in rodent studies. The A-ZIP/F mice, for instance, that feature a severe reduction in WAT, but only a partial reduction in BAT, due to the expression of dominant-negative protein ZIP/F, display reduced energy expenditure corrected by body weight and unaltered core body temperature in the fed state. Upon fasting, however, these mice display marked reductions in energy expenditure and core body temperature (torpor) [8,9]. Noteworthy, ZIP/F expression in this mouse model is driven by the aP2/FABP4 promoter, which is not specific to adipocytes, showing important activity in the brain, including the hypothalamus, and other tissues [10–13]. Similarly to A-ZIP/F, mice bearing a 50% reduction in WAT, but no changes in BAT mass, due to either whole-body or adipocyte-specific Bscl2/seipin deletion show unaltered and reduced energy expenditure expressed by body weight, respectively [14]. Finally, lipoatrophy induced by adipocyte deficiency of either mechanistic target of rapamycin complex 1 (mTORC1) or 2 (mTORC2) also does not alter mice energy expenditure [15,16].

Conversely to these studies showing no impact or a reduction in energy expenditure, partial lipoatrophy induced by whole-body deletion of early B-cell factor 1 (Ebf1) or a loss of function mutation in phosphoinositide 3-kinase (PI3K) (*Pikr3r1* Y657*), or by adipocyte-specific deletions of either the ubiquitin-conjugating enzyme E2 (Ube2i), or diacylglycerol acyltransferase 1 and 2 (DGAT1 and 2), is associated with increased energy expenditure [17–20]. Extending these findings, whole-body deletion of either 1-acylglycerol-3-phosphate-O-acyltransferase 2 (AGPAT2) or peroxisome proliferator activated receptor γ (PPARγ), which results in adipocyte death and complete absence of BAT and WAT, markedly increases energy expenditure at 23°C even after correction by lean mass [21,22]. Interestingly, in the latter, lipoatrophy-associated hypermetabolism was preserved at thermoneutrality (30°C), and associated with higher body temperature and metabolic inflexibility [22].

The reasons underlying these discrepancies in energy expenditure results in lipoatrophic patients and mice are unknown, but may be related to the different degrees and types of lipoatrophy, whether BAT mass and function were altered or not, as well as the usage of distinct procedures to correct and express energy expenditure results [23]. To overcome some of these caveats, we evaluated herein in well-controlled environmental conditions, energy balance and body temperature in severely lipoatrophic mice featuring complete absence of any visible WAT and BAT due to adipocyte-specific PPARγ deletion induced by adiponectin-Cre [24,25]. Noteworthy, the adiponectin promoter drives Cre expression, PPARγ deletion and death of mature brown and white adipocytes, generating a robust and specific mouse model of severe lipoatrophy [24–26].

## 2. Materials and Methods

### 2.1 Mice

Mice experimental procedures were approved by the Animal Care Committee of the Institute of Biomedical Sciences, University of Sao Paulo (#115/2016 and 6160250820, CEUA). All mice were male on a C57BL6/J background. PPARγ^Lox/Lox^ mice (B6.129-Ppargtm2Rev/j) were crossed with *adiponectin-cre* mice (B6; FVB-Tg(Adipoq-cre)1Evdr/J) as previously described [24,25] to obtain heterozygous PPARγ^Lox/WT^; adiponectin-cre^+/?^ offspring (WT refers to wildtype) in the F1 generation. These heterozygous mice were crossed with PPARγ^Lox/Lox^ mice to obtain PPARγ^Lox/Lox^; adiponectin-cre^+/−^ (referred henceforth as A-PPARγ KO mice) and their littermates PPARγ^Lox/Lox^ (referred henceforth as A-PPARγ WT). Mice genotypes were determined by PCR analysis of tail genomic DNA. 16 weeks old male mice were kept at either 17, or 23, or 30 ± 1°C on a 12:12 h light-dark cycle (lights on at 06:00 h) in environmental chambers (Promethion system, Sable System International), fed a nonpurified chow diet (70% carbohydrate, 20% protein, 10% fat, in %Kcal, NUVILAB CR-1®-Sogorb Inc., Sao Paulo, Brazil) and euthanized by exsanguination after anesthesia with isoflurane for tissue and blood harvesting. Body weight and food intake were measured weekly.

### 2.2. Treatments

In distinct dedicated cohorts, A-PPARγ WT and A-PPARγ KO mice were kept at 30 ± 1°C for at least one week and treated with either vehicle (1% DMSO in 0.2% carboxymethylcellulose) or the ryanodine receptor (RyR1) antagonist and calcium cycling inhibitor dantrolene (10 mg/kg/day, ip, Sigma) [27] or the acetyl-CoA carboxylase (ACC) inhibitor ND-630 (20 mg/kg/day, ip, InvivoChem) [28] or propranolol (0.75 g/L, Sigma) [29] in the drinking water for different periods and evaluated for energy expenditure as described below.

### 2.3. Energy balance

Mice were acclimated to the metabolic cages for 48 h and evaluated for energy expenditure (EE), oxygen consumption (VO_2_), carbon dioxide production (VCO_2_), spontaneous motor activity, respiratory exchange ratio (RER, VCO_2_/VO_2_ ratio), and food intake at different ambient temperatures during distinct periods in temperature-controlled cabinets (Promethion System, Sable Systems International, Las Vegas, USA). Oxygen consumption and carbon dioxide production were measured every 30 seconds, while activity was measured every second. Energy expenditure was directly calculated by Promethion software using the Weir equation [30].

### 2.4. Thermoneutral zone

In one dedicated protocol, mice acclimated at 30°C were evaluated for energy expenditure in Promethion System (Sable International) at different ambient temperatures during 4 h between 08:00 and 16:00 h in 4 consecutive days. The following order was applied: day 1-36°C from 08:00 to 12:00 h and 33°C from 12:00 to 16:00 h; day 2-30°C from 08:00 to 12:00 h and 27°C from 12:00 to 16:00 h; day 3-24°C from 08:00 to 12:00 h and 21°C from 12:00 to 16:00 h; day 4-18°C from 08:00 to 12:00 h and 15°C from 12:00 to 16:00 h. From 16:00 until 08:00 h, which comprises the dark period, mice were maintained at 30°C. The average energy expenditure from the last hour at each temperature was plotted to establish the thermoneutral zone, minimizing possible carryover effects.

### 2.5. Core body temperature

Mice were anesthetized with isoflurane and placed on a heat pad at 37°C for the insertion of a temperature probe transmitter (Sable Systems International) in the peritoneal cavity. The abdominal wall was sutured and the skin closed with surgical glue (Dermabond Topical Skin Adhesive; Johnson & Johnson, Brazil). After surgery, mice were treated with antibiotic (enrofloxacin 5 mg/kg sc) and analgesic (ketoprofen 5 mg/kg sc), and allowed to recover for 7 days at 30°C. After recovery, mice were adapted to the metabolic cages for 48 h at 30°C and evaluated for energy expenditure and core body temperature for 48 h consecutively at each of the following ambient temperatures: 30, 23 and 17°C. Core body temperature was evaluated by telemetry simultaneously with energy expenditure in the Promethion System.

### 2.6. RNA extraction and qPCR

Total RNA was extracted from gastrocnemius skeletal muscle, hypothalamus and liver, reverse transcribed, and destined for quantitative PCR analysis as previously described [31]. Primer nucleotide sequences are depicted in Table 1. Analysis of real-time PCR results was performed using the 2^-ΔΔC^_T_ method. Data are expressed as the ratio between the expression of the target gene and the housekeeping gene 36B4 (Rplp0), the expression of which was not significantly affected by mouse genotype.

**Table 1.**
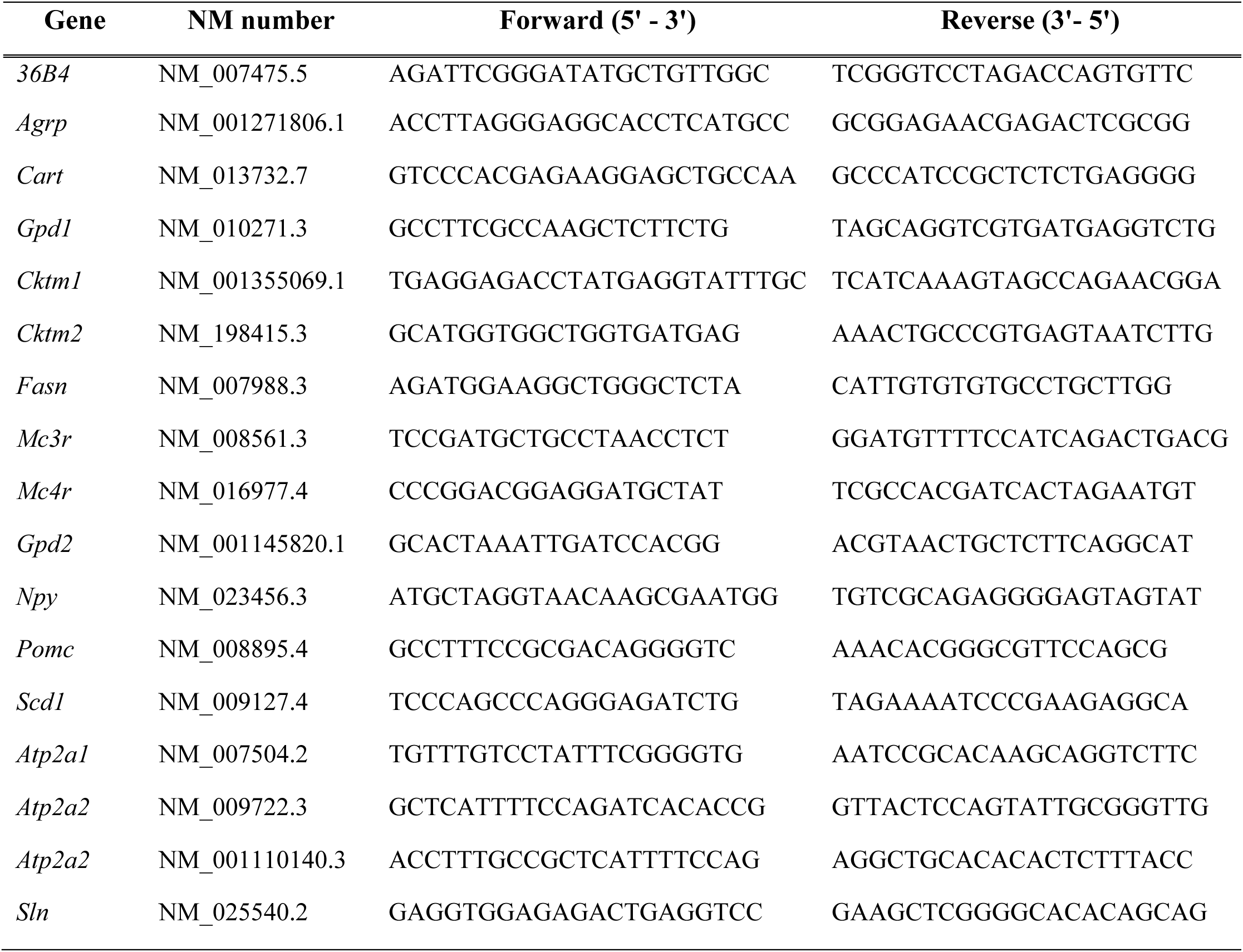
Primers used in qPCR.

### 2.7. Gas chromatography analysis

Liver fatty acid profile was measured by gas chromatography essentialy as previously described [24].

### 2.8. Western blotting

Liver was homogenized in buffer containing in mM, 50 HEPES, 40 NaCl, 50 NaF, 2 EDTA, 10 sodium pyrophosphate, 10 sodium glycerophosphate, 2 sodium orthovanadate, 1% Triton-X 100, and EDTA-free protease inhibitors and centrifuged at 13,000 rpm for 15 min at 4°C. Approximately, 20 μg of protein were separated on a gel, transferred to a PVDF membrane and immunoblotted with the following primary antibodies against ACC, FAS, SCD1, CPT1a, and Akt (#3662, 3189, 2438, 12252, and 9272, Cell Signaling). Membranes were washed, incubated with a secondary antibody (1:5000 in 5% nonfat milk in 0.05% TBS-Tween) and revealed with the chemiluminescent substrate ECL. Band densitometry was obtained with Image J software (National Institutes of Health, USA). Total Akt, whose content did not differ between genotypes and treatments, was used as a loading control.

### 2.9. Serum parameters

Serum from A-PPARγ WT and A-PPARγ KO mice kept at 30 ± 1°C and fasted for 5 h was evaluated for glucose, triacylglycerol and cholesterol using colorimetric assays (Labtest, Lagoa Santa, Brazil), as well as catecholamines (Biovision, US), free triiodothyronine (T3, ALPCO, US) and thyroid-stimulating hormone (TSH, Elabscience, US) by ELISA.

### 2.10. De novo fatty acid synthesis in liver slices

Liver slices of 1000 μm produced with a McIlwain Tissue Chopper were incubated in hermetically-closed vials, containing 1.5 mL of Krebs-Ringer Bicarbonate buffer pH 7.4 composed of (in mM) 5 glucose, 0.51 MgCl_2_, 4.56 KCl, 119.8 NaCl, 0.7 Na_2_HPO_4_, 1.3 NaH_2_PO_4_, 15.0 NaHCO_3_, 1% essentially fatty acid-free albumin, containing 2 mM acetate and 0.2 μCi/vial [1-^14^C] acetate for evaluation of fatty acid synthesis (*de novo* lipogenesis). After 2 h at 37°C, the medium was acidified and liver slices were washed with saline and destined for lipid extraction with chloroform/methanol for measurement of acetate incorporation into triacylglycerol-fatty acids as described [32,33]. Values were expressed per mg of protein to correct for cell number.

### 2.11. Statistical analysis

Data are expressed as mean ± SEM. Student’s *t*-test was used to test the impact of mouse genotype (A-PPARγ WT vs A-PPARγ KO) at each specific ambient temperature. One-way ANOVA followed by Tukey’s post hoc test were used to evaluate the impact of different ambient temperatures (17, 23 and 30°C) on either A-PPARγ WT or A-PPARγ KO mice. ANOVA for repeated measures was used to compare energy expenditure before and after propranolol treatment. Multifactorial ANOVA followed by Tukey’s post hoc test was used to evaluate the impact of treatments (eitherdantrolene or ND-630), genotypes and their interactions. Student’s *t*-test and ANOVA were performed using GraphPad Prism 10.6. ANCOVA was performed using CalR (https://calrapp.org/) to evaluate the impact of body weight and lean mass on energy expenditure. The significance level was set at *p* ≤ 0.05.

## 3. Results

As illustrated in Figure 1, despite the complete absence of WAT and BAT, 16-weeks old male A-PPARγ KO featured at 30°C augmented body weight (A), lean (B), fat (C), liver (D), stomach (E), intestine (F), pancreas (G), spleen (H), heart (I) and adrenal gland (J), but reduced gastrocnemius skeletal muscle (K) masses when compared to their littermate A-PPARγ WT. Furthermore, A-PPARγ KO at 30°C showed increased fasting serum glucose, triacylglycerol and cholesterol when compared to A-PPARγ WT (Table 2).

**Figure 1.**
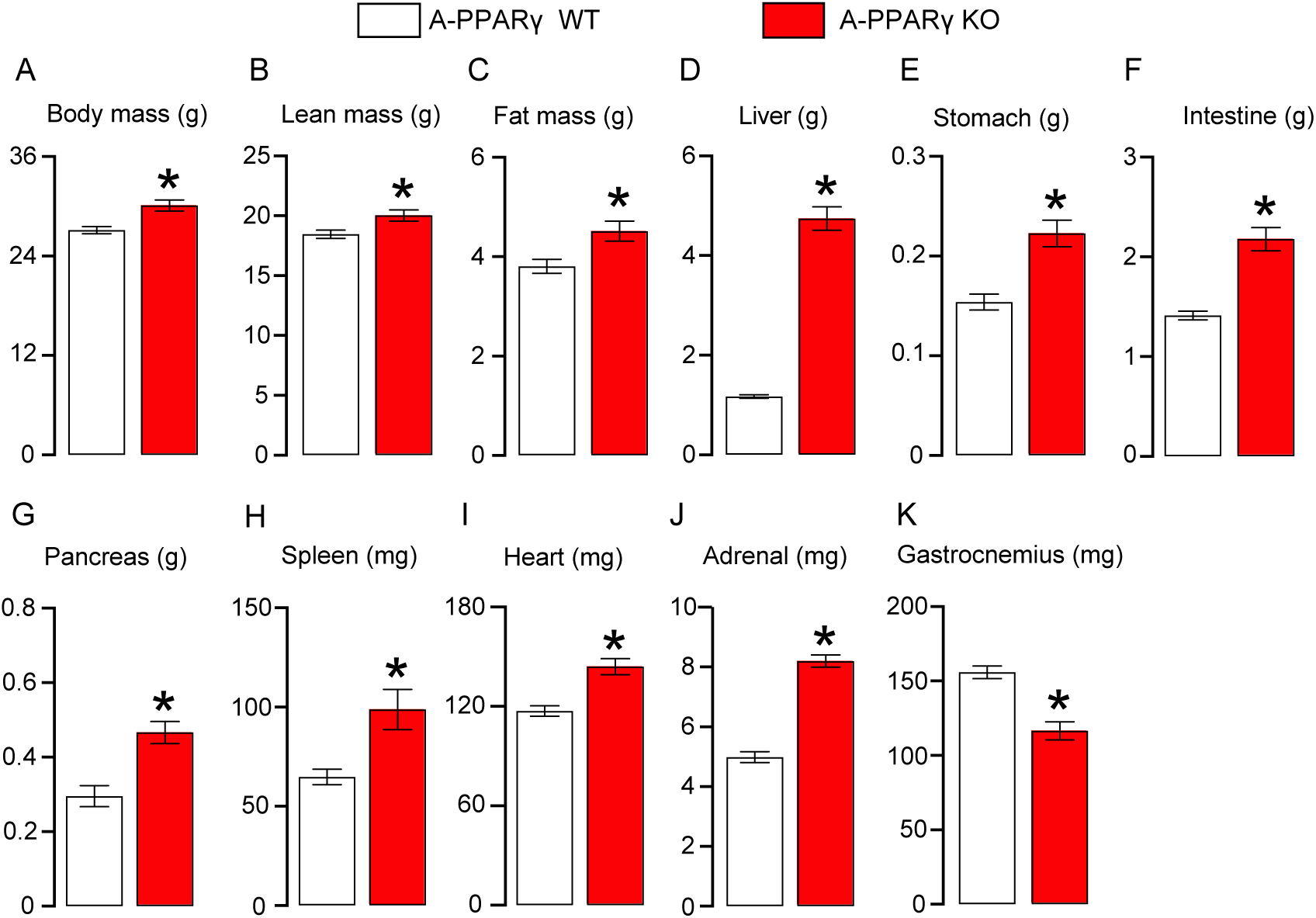
Severe lipoatrophy increases body weight, lean and fat masses, promotes organomegaly, but reduces skeletal muscle mass in mice at 30°C. Body (A), lean (B), fat (C), liver (D), stomach (E), intestine (empty small + large intestine, F), pancreas (G), spleen (H), heart (I), adrenal gland (J), and gastrocnemius skeletal muscle (K) masses in 16 weeks old littermates PPARγ flox (A-PPARγ WT) and PPARγ flox adiponectin-Cre^+^ (A-PPARγ KO) male mice kept at 30°C. Results are expressed as mean ± SEM. Unpaired Student’s *t*-test, * *p* ≤ 0.05.

**Table 2.** Serum parameters in A-PPARγ WT and KO mice kept at 30°C.

| | A-PPARγ WT | A-PPARγ KO |
| --- | --- | --- |
| Glucose (mg/ dL) | 166.6 ± 7.9 | 251.9 ± 13.4* |
| TAG (mg/ dL) | 72.2 ± 7.5 | 180.2 ± 22.4* |
| Cholesterol (mg/ dL) | 88.4 ± 2.4 | 176.2 ± 9.9* |

Results are mean ± SEM, n= 6-8 mice per group. \* p< 0.05 versus A-PPARγ WT.

Along with severe lipoatrophy and organomegaly, A-PPARγ KO displayed higher rates of energy expenditure at both the light and dark cycles at 30°C (Figure 2A), ensuing a higher cumulative energy expenditure over 5 days (Figure 2D). Similarly to 30°C, A-PPARγ KO also displayed higher rates of energy expenditure and cumulative energy expenditure than A-PPARγ WT at 23°C, such a difference that was more prominent at the dark period (Figure 2B and E). When exposed to 17°C, however, A-PPARγ KO displayed higher rates of energy expenditure than A-PPARγ WT only at the dark period in the first 72 h, and at both light and dark periods in the last 48 h (Figure 2C). As the result of this distinct behavior, the difference in cumulative energy expenditure between A-PPARγ KO and A-PPARγ WT was less robust than that seen at 30 and 23°C (Figure 2F). Re-organization and re-analysis of these results by genotype reveals that reducing ambient temperature from 30 to 23 and subsequently to 17°C, gradually increased energy expenditure in A-PPARγ WT at both light and dark cycles, with the highest rates seen at 17°C (Figure 2G and I). Differently from A-PPARγ WT, exposure of A-PPARγ KO to 23°C increased energy expenditure only at the dark cycle, while exposure to 17°C further increased energy expenditure at both the light and dark cycles to rates that were significantly higher than those found at 23 and 30°C (Figure 2H and J). Indeed, A-PPARγ KO showed at 17°C a higher cumulative energy expenditure than at 23 and 30°C, while a significant difference between 23 and 30°C was apparent only at later time points (Figure 2H and J). Altogether, these findings indicate that severe lipoatrophic mice are hypermetabolic at all ambient temperatures investigated, emphasizing the existence of recruitable WAT and BAT-independent mechanisms to increase energy expenditure.

**Figure 2.**
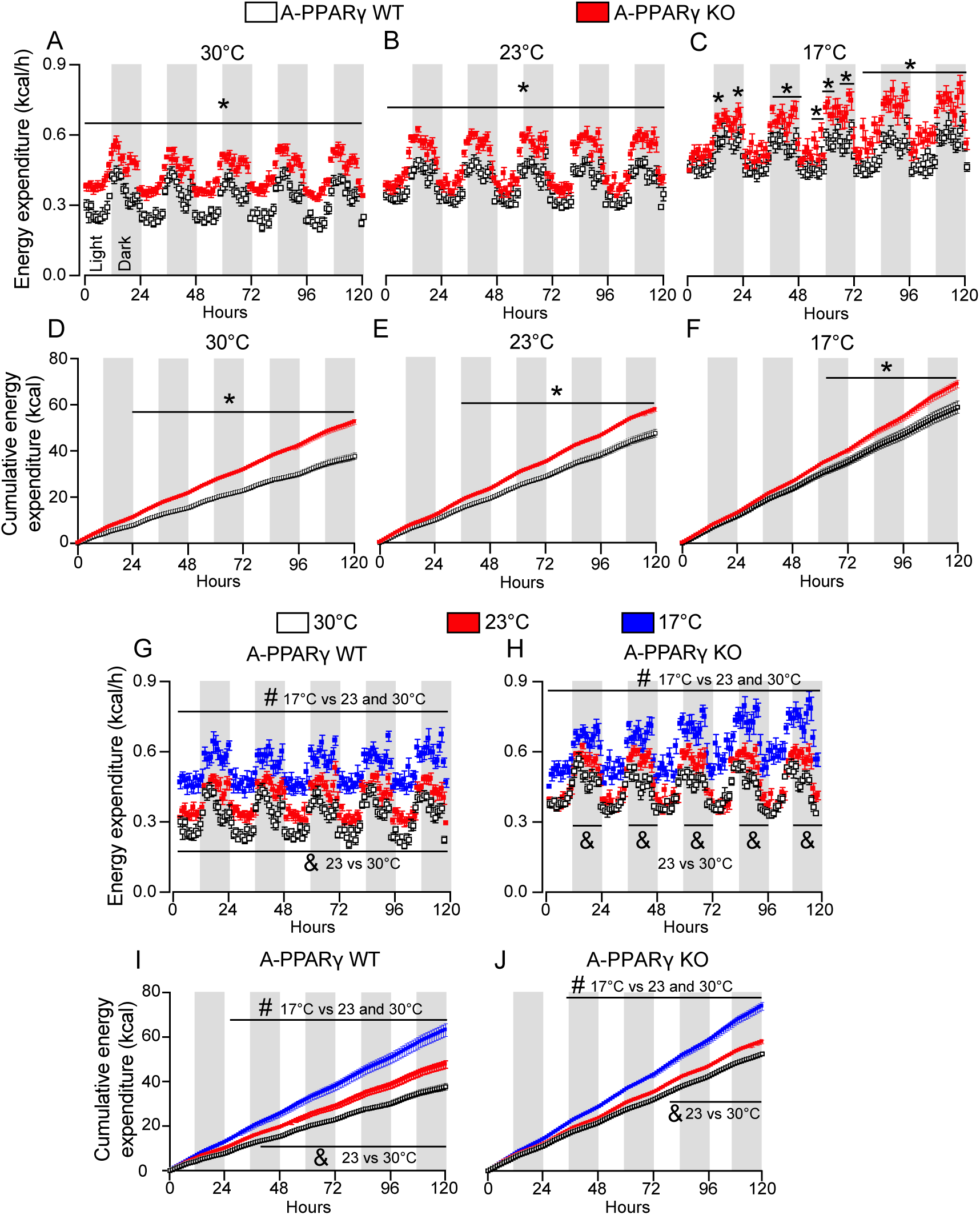
Severe lipoatrophy increases energy expenditure in mice at different ambient temperatures. Energy expenditure (A-C) and cumulative energy expenditure (D-F) in 16 weeks old, littermates PPARγ flox (A-PPARγ WT) and PPARγ flox adiponectin-Cre^+^ (A-PPARγ KO) male mice consecutively exposed to 30°C, 23°C and 17°C for 5 days at each temperature. In panels G-H and I-J, energy expenditure and cumulative energy expenditure results, respectively, were plotted by genotype. Results are expressed as mean ± SEM. Results displayed in A-F were analyzed by Unpaired Student’s *t*-test, * *p* ≤ 0.05, A-PPARγ WT vs A-PPARγ KO at each ambient temperature. In G to J, results were analyzed by One-way ANOVA followed by Tukey post hoc test. # *p* ≤ 0.05, 17°C vs 23°C and 30°C; & *p* ≤ 0.05, 23°C vs 30°C.

As illustrated in Figure 3, reducing ambient temperature to 23 (Fig. 3B) and 17°C (Fig. 3C) enhances respiratory exchange rate (RER) circadian variation in both A-PPARγ WT and KO, as evidenced by the lower (nadir) and higher (peak) RER values at the light and dark cycles, respectively, when compared to 30°C (Fig. 3A). Interestingly, there were no significant differences in RER between A-PPARγ KO and WT at 30°C (Fig. 3A), while at 23 and 17°C (Fig. 3B and C), A-PPARγ KO displayed a narrower circadian variation in RER than A-PPARγ WT characterized by higher and lower RER values at the light and dark cycles, respectively, a phenotype generally defined as metabolic inflexibility. As depicted in Figure 3D-F, A-PPARγ KO displayed higher cumulative food intake than A-PPARγ WT at all ambient temperatures investigated. Plotting these results by genotype indicates that reducing ambient temperature significantly increases cumulative food intake in A-PPARγ WT, but not A-PPARγ KO (Figure 3G and H). In contrast to food intake, A-PPARγ KO displayed reduced ambulatory activity when compared to A-PPARγ WT, as evidenced by the reduced total distance traveled at all ambient temperatures (Figure 3I-K). Plotting the results by genotype reveals that reducing ambient temperature to 17°C significantly reduced total distance traveled by both A-PPARγ WT and KO (Figure 3L and M). In spite of a significant reduction in A-PPARγ KO (Figure 3M), there were, however, no differences in total distance traveled by A-PPARγ WT between 30 and 23°C (Figure 3L). Altogether, these findings indicate that A-PPARγ KO mice feature metabolic inflexibility below thermoneutrality, along an increased, but non-adjustable food intake, and a reduced, but adjustable ambulatory activity at different ambient temperatures.

**Figure 3.**
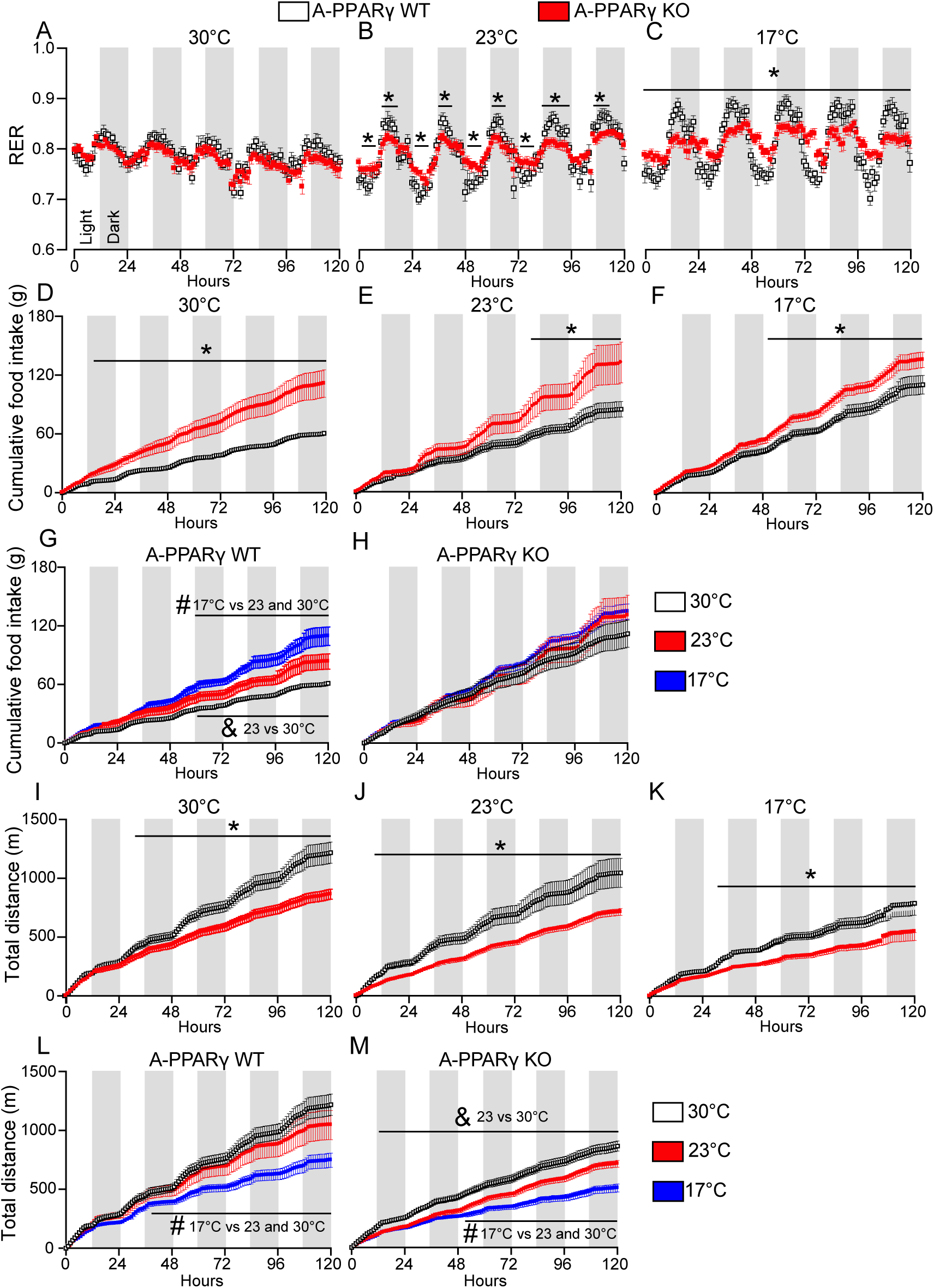
Severe lipoatrophy promotes metabolic inflexibility, increases food intake and reduces total distance traveled. Respiratory exchange ratio (RER, A-C), cumulative food intake (D-F), and total distance traveled (I-K) in 16 weeks old, littermate PPARγ flox (A-PPARγ WT) and PPARγ flox adiponectin-Cre^+^ (A-PPARγ KO) male mice consecutively exposed to 30°C, 23°C and 17°C for 5 days at each temperature. In panels G-H and L-M, cumulative food intake and total distance traveled, respectively, were plotted by genotype. Results are expressed as mean ± SEM. Results displayed in A-F and I-J were analyzed by Unpaired Student’s *t*-test, * *p* ≤ 0.05, A-PPARγ WT vs A-PPARγ KO at each ambient temperature. In G-H and L-M, results were analyzed by One-way ANOVA followed by Tukey post hoc test. # *p* ≤ 0.05, 17°C vs 23°C and 30°C; & *p* ≤ 0.05, 23°C vs 30°C.

As A-PPARγ KO are heavier and display higher lean mass than A-PPARγ WT, we next evaluated the impact of body and lean masses on energy expenditure by performing an ANCOVA analysis. As illustrated in Figure 4A and B, there were no significant effects of body (p= 0.2967) and lean (p= 0.3286) masses on mice energy expenditure at 30°C. Furthermore, A-PPARγ KO display higher rates of energy expenditure at 30°C than A-PPARγ WT even after correction by lean mass (Figure 4C). Altogether, these findings indicate that the increase in energy expenditure displayed by A-PPARγ KO is not exclusively due to their higher body weight and lean mass. Next, we exposed A-PPARγ WT and KO to different ambient temperatures varying from 36 to 15°C (a 3°C reduction at every 4 h exclusively between 08:00 and 16:00 h at the light cycle) to evaluate their thermoneutral zone. As illustrated in Figure 4D, A-PPARγ KO displayed higher energy expenditure rates than A-PPARγ WT across all ambient temperatures tested, with the minimal rates (nadir) occurring between 24 and 33°C. A-PPARγ WT, on the other hand, displayed minimal energy expenditure rates between 30 and 33°C. These findings indicate a widening and repositioning of thermoneutral zone toward lower ambient temperatures in A-PPARγ KO.

**Figure 4.**
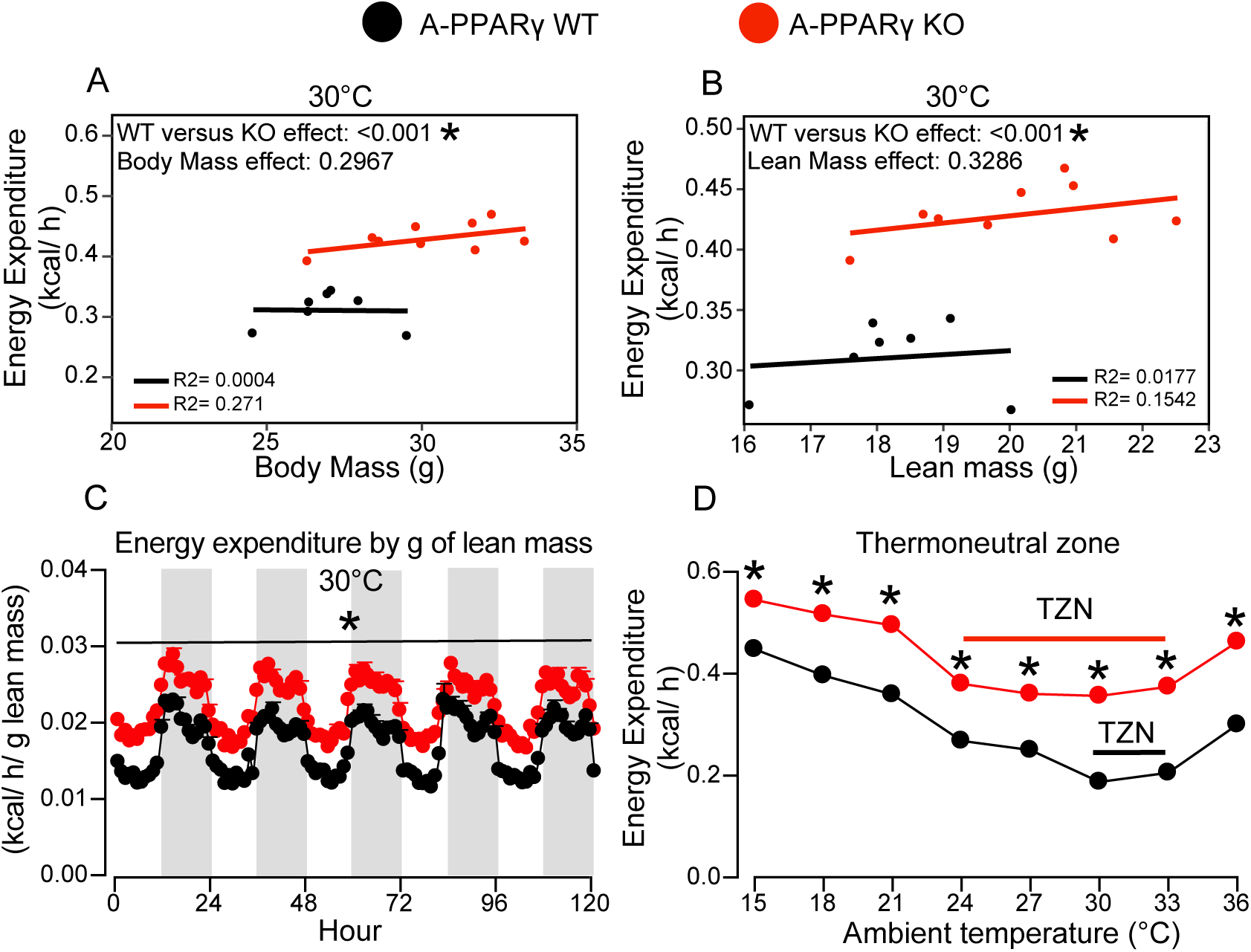
Hypermetabolism in severely lipoatrophic mice is not due to increased body and lean masses, and is associated with widening of the thermoneutral zone. ANCOVA analysis of energy expenditure using either body mass (A) or lean mass (B) as covariates perfomed using CalR, and energy expenditure expressed per gram of lean mass (C) of 16 weeks old, littermates PPARγ flox (A-PPARγ WT) and PPARγ flox adiponectin-Cre^+^ (A-PPARγ KO) male mice kept at 30°C for 14 days. In Panel C, results expressed as mean ± SEM were analyzed by Unpaired Student’s *t*-test, * *p* ≤ 0.05, A-PPARγ WT vs A-PPARγ KO, n=7-9 mice per genotype. Panel D depicts rates of energy expenditure of a different cohort of 16 weeks old, littermate PPARγ flox (A-PPARγ WT) and PPARγ flox adiponectin-Cre^+^ (A-PPARγ KO) male mice exposed to 36, 33, 30, 27, 24, 21, 18 and 15°C during 4 h in the light period from 08:00 to 16:00 h during 4 consecutive days. Plotted results are the average of the last hour at each temperature. Unpaired Student’s *t*-test, * *p* ≤ 0.05, A-PPARγ WT vs A-PPARγ KO, n=6-10 mice per genotype.

Subsequently, we implanted thermoprobes in the peritoneal cavity of a new mice cohort to investigate the impact of different ambient temperatures on energy expenditure and core body temperature simultaneously. At 30°C, A-PPARγ KO enhanced rates of energy expenditure were associated with a high core body temperature (hyperthermia) when compared to A-PPARγ WT at both the light and dark cycles (Figure 5A and B). At 23°C, however, A-PPARγ KO also displayed enhanced energy expenditure rates, but this was associated with no significant changes in core body temperature (Figure 5C and D). Interestingly, some, but not all A-PPARγ KO mice featured a bout of hypothermia at the beginning of the light cycle, which did not achieve statistical significance. Upon exposure to 17°C, however, A-PPARγ KO also displayed periods of higher energy expenditure at both the light and dark cycles in comparison to A-PPARγ WT, which occurred alongside with bouts of hypothermia that were more prominent at the beginning of the light cycle (Figure 5E and F). These findings indicate that A-PPARγ KO are hyperthermic at thermoneutrality, but have difficulties preserving internal heat and maintaining core body temperature upon exposure to ambient temperatures below thermoneutrality.

**Figure 5.**
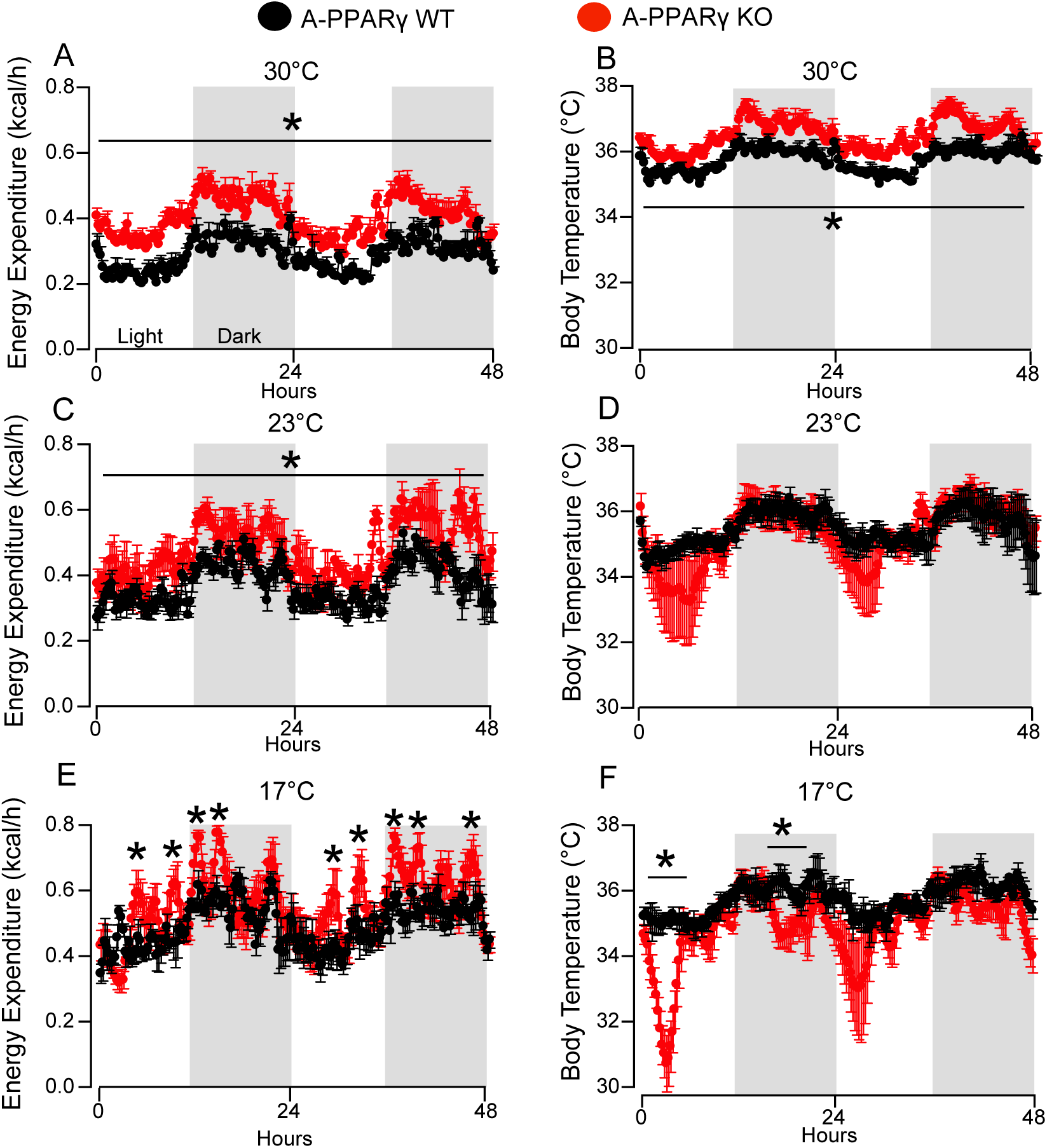
Severe lipoatrophy is associated with hyperthermia at 30°C and bouts of hypothermia at 17°C. Energy expenditure (A, C and E) and core body temperature (B, D and F) in 16 weeks old, littermate PPARγ flox (A-PPARγ WT) and PPARγ flox adiponectin-Cre^+^ (A-PPARγ KO) male mice consecutively exposed to 30°C, 23°C and 17°C for 48 h at each temperature. Unpaired Student’s *t*-test, * *p* ≤ 0.05, A-PPARγ WT vs A-PPARγ KO, n= 8 mice per genotype.

In an attempt to investigate the mechanisms underlying A-PPARγ KO enhanced energy expenditure at 30°C, we performed a gene expression analysis of hypothalamus, the major regulator of energy balance. As illustrated in Figure 6A, A-PPARγ KO kept at 30°C displayed elevated hypothalamic mRNA content of anorexigenic and pro-energy expenditure proopiomelanocortin (POMC), along with increased content of the orexigenic agouti-related protein (AgRP) and neuropeptide Y (NPY). Considering that elevated hypothalamic POMC levels are associated with enhanced energy expenditure and sympathetic activity (reviewed in [34]), we next administered propranolol to mice to investigate a possible implication of β-adrenergic receptor signaling as mediator of the hypermetabolism featured by lipoatrophic mice at 30°C. As illustrated in Figure 6B, propranolol treatment (0.75 g/L in drinking water) significantly reduced energy expenditure in both A-PPARγ WT and KO mice, but it did not abolish the difference in energy expenditure between genotypes, indicating that β-adrenergic receptor signaling is not a major system driving hypermetabolism in lipoatrophic mice. Further supporting these findings, there were no changes in serum catecholamines, as well as free triiodothyronine (T3) and thyroid-stimulating hormones (TSH) levels between genotypes (Table 3), excluding a participation of sympathetic nervous system and thyroid hormones as mediators of lipoatrophic mice hypermetabolism. Considering that skeletal muscle is an important determinant of energy expenditure, we next evaluated the gene expression profile of thermogenesis-related proteins in gastrocnemius skeletal muscle. As illustrated in Figure 6C, A-PPARγ KO at 30°C showed a marked increase in gastrocnemius skeletal muscle mRNA content of proteins that compose the energy-expending calcium futile cycle namely sarcoplasmic/endoplasmic reticulum Ca^2+^-ATPases 2a and 2b (SERCA2a and 2b, respectively) and sarcolipin (SLN), a protein that reduces SERCA affinity for calcium, reducing and uncoupling pumping activity from ATP consumption [35,36]. To test the contribution of SERCA-SLN mediated calcium futile cycle to the increased energy expenditure featured by lipoatrophic mice, we next treated A-PPARγ WT and KO at 30°C with dantrolene, a ryanodine receptor (RyR1) antagonist and calcium cycling inhibitor. As illustrated in Figure 6D, dantrolene administration did not significantly affect rates of energy expenditure in both A-PPARγ WT and KO mice, excluding an involvement of calcium cycling as mediator of the higher energy expenditure featured by lipoatrophic mice.

**Figure 6.**
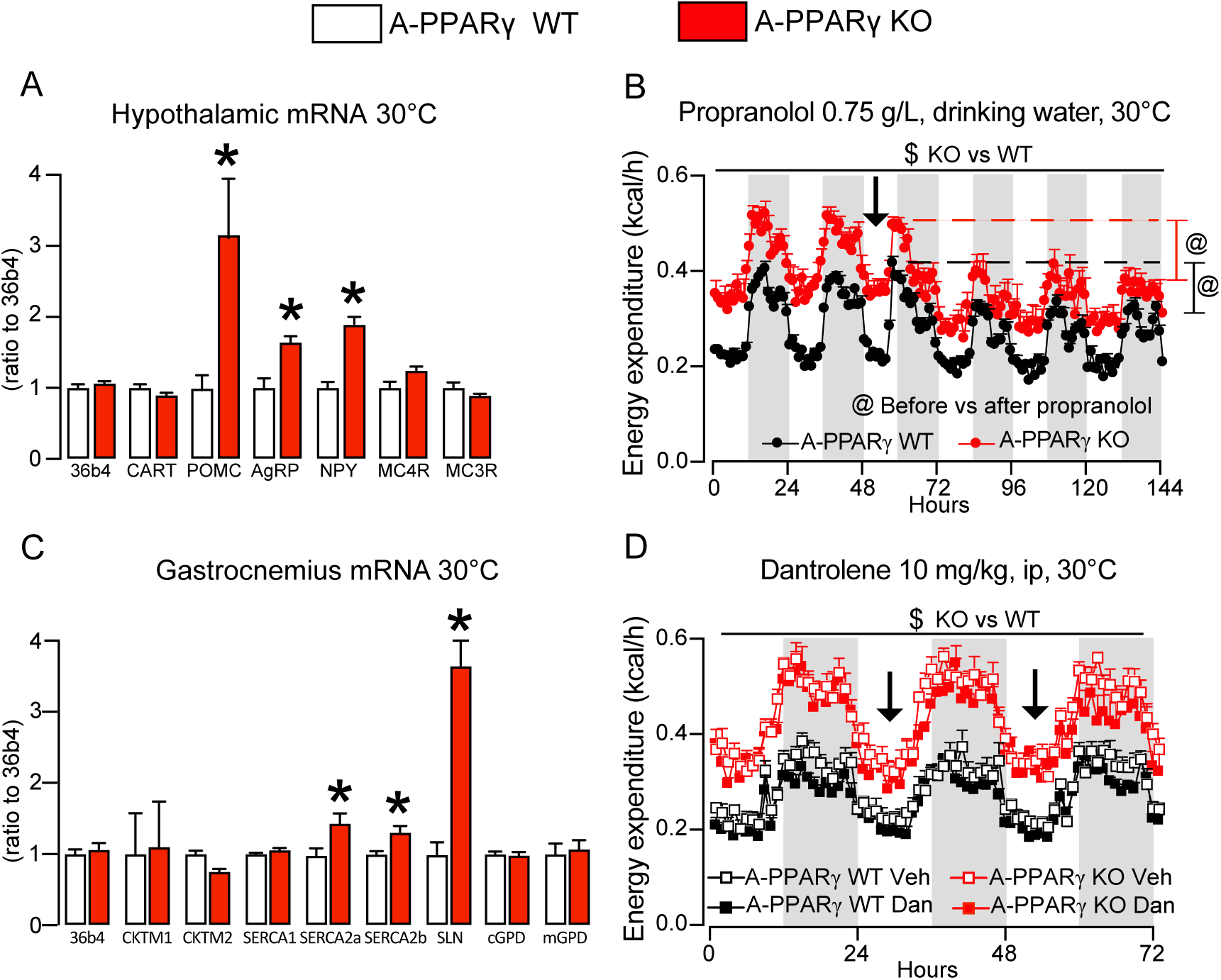
Hypermetabolism in severely lipoatrophic mice is not affected by β-adrenergic receptor signaling or calcium cycling pharmacological blockade. Hypothalamus (A) and gastrocnemius skeletal muscle (C) gene expression analysis in 16 weeks old, littermate PPARγ flox (A-PPARγ WT) and PPARγ flox adiponectin-Cre^+^ (A-PPARγ KO) male mice kept at 30°C. Unpaired Student’s *t*-test, * *p* ≤ 0.05, A-PPARγ WT vs A-PPARγ KO, n= 8 mice per genotype. Rates of energy expenditure in 2 different cohorts of 16 weeks old, littermate PPARγ flox (A-PPARγ WT) and PPARγ flox adiponectin-Cre^+^ (A-PPARγ KO) male mice kept at 30°C. One cohort was treated with propranolol (0.75 g/L during 4 days) in the drinking water (Panel B), while the other was treated with daily injections of either dantrolene (10 mg/kg/day, ip, during 2 days) or vehicle (1% DMSO in 0.2% carboxymethylcellulose) (Panel D). $ p < 0.05 A-PPARγ KO vs WT, Unpaired Student’s *t*-test and ANOVA; @ p < 0.05 same genotype before vs after propranolol, ANOVA for repeated measures.

**Table 3.** Serum hormones in A-PPARγ WT and KO mice kept at 30°C.

| | A-PPARγ WT | A-PPARγ KO |
| --- | --- | --- |
| Catecholamines (pg/ mL) | 29.5 ± 0.69 | 28.3 ± 0.60 |
| Free T3 (pg/ mL) | 1.35 ± 0.20 | 1.65 ± 0.08 |
| TSH (ng/ mL) | 14.6 ± 0.90 | 16.9 ± 0.91 |

Results are mean ± SEM, n= 6-8 mice per group. \* p< 0.05 versus A-PPARγ WT.

As depicted in Figure 7, analysis of gene expression profile in A-PPARγ KO liver revealed significant increases in mRNA content of SERCA2a; cytosolic and mitochondrial glycerol 3-phosphate dehydrogenase (cGPD and mGPD, respectively) that catalyze a futile metabolic cycle involving the interconversion between glycerol 3-phosphate and dihydroxyacetone 3-phosphate; and fatty acid synthase (FAS) and stearoyl-CoA desaturase (SCD1) involved in *de novo* lipogenesis (Figure 7A). Analysis of liver fatty acid profile by gas chromatography showed not only an increase in the content of *de novo* lipogenesis products palmitic (16:0), palmitoleic (16:1n7) and oleic (18:1) acids, but also a significant reduction in linoleic (18:2n6) and alpha-linolenic (18:3n3) acids, which are preferentially β-oxidized [24,25,37–39] (Figure 7B). Confirming the upregulation of *de novo* lipogenesis, A-PPARγ KO liver slices had higher rates of acetate incorporation into triacylglycerol-fatty acids than liver slices of A-PPARγ WT (Figure 7C). Then, to test the contribution of *de novo* lipogenesis to lipoatrophic mice hypermetabolism, we treated A-PPARγ WT and KO at 30°C with ND-630, a potent pharmacological inhibitor of acetyl-CoA carboxylase (ACC) [40]. We chose to inhibit ACC instead of FAS to avoid malonyl-CoA accumulation, allosteric CPT1 inhibition and an impairment of mitochondrial fatty acid β-oxidation. As illustrated in Figure 7D, A-PPARγ KO treated with vehicle showed increased liver content of ACC, FAS, SCD1 and a trend for an increase in carnitine palmitoyl transferase 1 (CPT1) when compared to vehicle treated A-PPARγ WT, such differences that were fully abolished by ACC inhibition with ND-630. Along with the reduction in *de novo* lipogenesis enzymes and CPT1, ND-630 also reduced energy expenditure and RER values, indicating increased β-oxidation, in both A-PPARγ WT and KO mice (Figure 7E and F). Noteworthy, ND-630 did not significantly affect A-PPARγ WT and KO food intake (Figure 7G). Altogether, these findings indicate that liver *de novo* lipogenesis partly contributes to the increased energy expenditure and hypermetabolism displayed by lipoatrophic A-PPARγ KO mice at 30°C.

**Figure 7.**
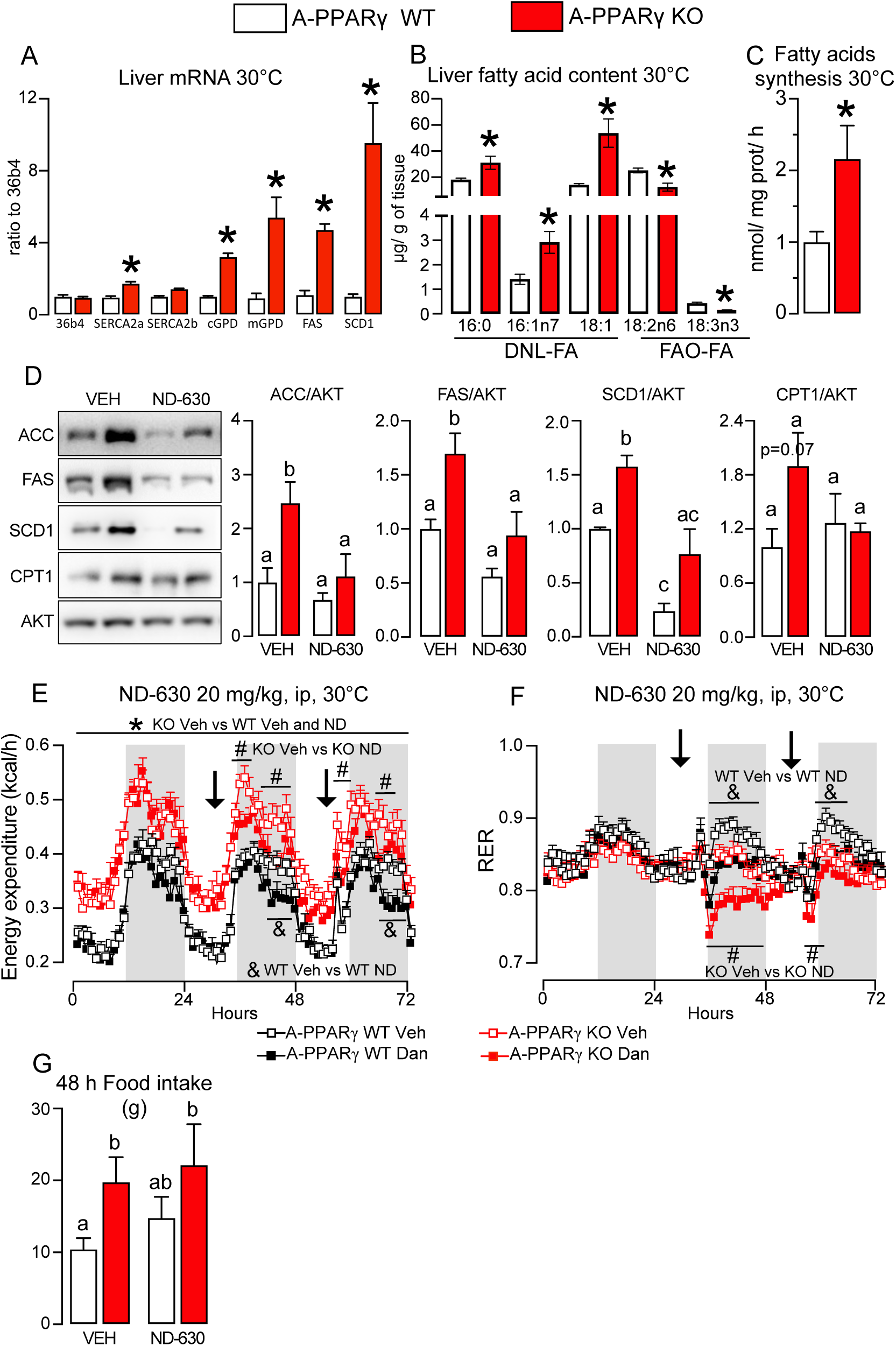
Hypermetabolism in severely lipoatrophic mice is partly attenuated by pharmacological inhibition of ACC and *de novo* fatty acid synthesis. Liver gene expression profile (A), fatty acid content (B), and fatty acid synthesis (C, labeled acetate incorporation into triacylglycerol-fatty acids) in 16 weeks old, littermate PPARγ flox (A-PPARγ WT) and PPARγ flox adiponectin-Cre^+^ (A-PPARγ KO) male mice kept at 30°C. Unpaired Student’s *t*-test, * *p* ≤ 0.05, A-PPARγ WT vs A-PPARγ KO, n= 8 mice per genotype. Liver protein content of *de novo* lipogenesis enzymes and carnitine palmitoyl transferase 1a (CPT1a), and rates of energy expenditure (E), respiratory exchange rate (RER, F) and food intake (G) in 16 weeks old, littermate PPARγ flox (A-PPARγ WT) and PPARγ flox adiponectin-Cre^+^ (A-PPARγ KO) male mice kept at 30°C treated with the acetyl-CoA carboxylase (ACC) inhibitor ND-630 (20 mg/kg/day during 2 days) or vehicle (1% DMSO in 0.2% carboxymethylcellulose). Two-way ANOVA, * A-PPARγ KO Veh vs. WT treated with Veh or ND-630; # A-PPARγ KO Veh vs. KO treated with ND-630; & A-PPARγ WT Veh vs. WT treated with ND-630. n= 8 mice per group. In D and G, means not sharing a common superscript letter are significantly different, Two-way ANOVA.

## 4. Discussion

We investigated herein how severe lipoatrophy impacts energy balance at different ambient temperatures profiting from a specific and robust mouse model, in which, mature brown and white adipocyte death is induced by PPARγ deletion driven by adiponectin-Cre. Severely lipoatrophic mice are heavier, hypermetabolic and hyperphagic, and feature a widened thermoneutral zone, a markedly reduced ambulatory activity, a prominent metabolic inflexibility at both 23 and 17°C, and an unstable thermal behavior characterized by hyperthermia at 30°C, normothermia at 23°C, and bouts of hypothermia at 17°C. Notably, hypermetabolism in lipoatrophic mice at 30°C is not driven by thyroid hormones, impaired insulation or the increased body and lean masses and is not affected by pharmacological blockade of either β-adrenergic receptor signaling with propranolol or skeletal muscle SERCA-SLN-mediated calcium cycling with dantrolene, but is partially attenuated by pharmacological inhibition of ACC and *de novo* lipogenesis with ND-630.

Severely lipoatrophic mice featured higher rates of energy expenditure than littermate controls at all ambient temperatures investigated. Such phenotype was associated with lower ambulatory activity and a marked hyperphagia, altogether indicating an increased contribution of the thermic effect of food-in detriment of activity-related energy expenditure to lipoatrophic mice hypermetabolism. Considering that lipoatrophic mice do not have BAT and thus facultative diet-induced thermogenesis, only the obligatory component of the thermic effect of food, i.e., the energy used to digest and absorb nutrients, is expected to be enhanced in lipoatrophic mice. Interestingly, in contrast to the gradual increase in food intake shown by controls upon exposure to 23 and 17°C, hyperphagia in lipoatrophic mice, which is caused by the absence of leptin and is associated with high hypothalamic mRNA expression of orexigenic peptides NPY and AgRP, was not further enhanced by exposure to 23 and 17°C, despite the marked increase in energy expenditure seen at these ambient temperatures. Such dissociation between food intake and energy expenditure may be due to the magnitude of hyperphagia attained in lipoatrophic mice, which may be high enough to accommodate the increase in energy requirement induced by exposure to lower ambient temperatures.

Altogether, the dissociation between food intake and energy expenditure, the absence of diet-induced thermogenesis, along with the marked reduction in ambulatory activity suggest an important contribution of basal metabolic rate, i. e., the energy cost of all processes that sustain life including homeothermy, to hypermetabolism in lipoatrophic mice. Considering the impaired insulation associated with almost complete WAT absence, one putative cause of hypermetabolism in lipoatrophic mice may be a compensatory activation of thermogenic processes to maintain homeothermy in a condition of increased heat loss to the environment. Although this may be the case at 17°C, as evidenced by the impaired heat conservation and bouts of hypothermia, at 23 and 30°C hypermetabolism in lipoatrophic mice was associated with normothermia and hyperthermia, respectively, excluding a role of poor insulation as the underlying cause of the enhanced energy expenditure featured by lipoatrophic mice at these ambient temperatures. Similarly to the impaired insulation, ANCOVA analysis revealed a minor, nonsignificant contribution of increased body and lean masses to hypermetabolism in lipoatrophic mice.

Interestingly, lipoatrophic mice hypermetabolism is associated with widening and repositioning of thermoneutral zone characterized by energy expenditure rates nadir occurring between 24 and 33°C in comparison to 30 and 33°C in controls. Thermoneutral zone refers to a range of ambient temperatures in which homeothermy is maintained by passive heat exchange with the environment and thermogenic processes are at their minimal activity. Noteworthy, although thyroid hormones are major enhancers of basal metabolic rates [41,42] and hyperthyroidism is associated with widening of the thermoneutral zone and pyrexia (defended increased body temperature) even in UCP1-deficient mice [43], the absence of significant changes in serum free T3 and TSH levels excludes a possible role of thyroid hormones as drivers of lipoatrophic mice hypermetabolism at 30°C. Along with thyroid hormones, serum catecholamines were not altered and β-adrenergic receptor signaling blockade with propranolol similarly reduced energy expenditure in both controls and lipoatrophic mice, excluding a likely involvement of the sympathetic nervous system as an inducer of lipoatrophic mice hypermetabolism. Furthermore, although gene expression analysis of gastrocnemius muscle revealed a significant increase in mRNA content of futile calcium cycling components SLN and SERCAs, inhibition of ryanodine receptor (RyR1)-mediated calcium cycling with dantrolene did not alter lipoatrophic mice energy expenditure at 30°C, also excluding a role of muscle calcium cycling as a driver of lipoatrophic mice hypermetabolism.

Extending previous findings [24,25], lipoatrophy was associated at 30°C with a marked increase in liver *de novo* fatty acid synthesis, as evidenced by the enhanced hepatic rates of acetate incorporation into fatty acids and liver contents of *de novo* lipogenesis enzymes ACC, FAS and SCD1 and products namely palmitic (16:0), palmitoleic (16:1n7) and oleic (18:1) acids. Interestingly, pharmacological inhibition of ACC and *de novo* lipogenesis with ND-630 not only reduced energy expenditure in control mice, but partially abolished the hypermetabolism of lipoatrophic mice, such effects that were more evident at the dark period when rates of *de novo* fatty acid synthesis are at their highest [44]. Altogether, these findings indicate that enhanced liver *de novo* lipogenesis is an important contributor to lipoatrophic mice hypermetabolism, a finding that is consistent not only with the high energy requirement for *de novo* palmitic acid synthesis (8 ATP and 14 NADPH per molecule), but also with the 5-fold increase in liver mass in lipoatrophic mice. Noteworthy, ACC inhibition also significantly reduced RER in both genotypes indicating an increase in β-oxidation, most likely as the result of an alleviation of CPT1 inhibition by the reduction in ACC product malonyl-CoA. Interestingly, ACC inhibition also reduced its liver content and those of FAS and SCD1, such findings that are consistent with the previously identified positive feedback loop in which *de novo* lipogenesis product palmitoleic acid activates PPARα that, in turn, enhances the transcription of ACC, FAS and SCD1 and *de novo* lipogenesis as evaluated *in vivo* by the incorporation of tritiated water and labeled acetate into triacylglycerol-fatty acids [45,46].

Aside from the mechanisms driving lipoatrophic mice hypermetabolism, which may involve processes not investigated in this study such as the GPD1 and GPD2-mediated futile cycle, among others, one interesting question that emerges relates to why the cost of living without mature adipocytes and extremely limited energy stores is higher. Severely lipoatrophic mice rely almost exclusively on liver glycogen and triacylglycerol as energy stores, and when fasted, they enter torpor much sooner than controls as a strategy to reduce energy expenditure in face of limited energy availability [9]. Considering that the absence of WAT and thus of an organ for safe energy storage impose a severe metabolic distress characterized by lipotoxicity, inflammation, insulin resistance, hyperglycemia and hyperinsulinemia, lipoatrophic mice hypermetabolism may be the energetic burden required to adapt and survive in this chronic stressful metabolic environment. Indeed, severe lipoatrophy/lipodystrophy could be classified as a state of allostatic overload considering its main features namely hypermetabolism, high serum levels of stress-related hormones such as glucocorticoids, fibroblast growth factor 21 (FGF21), growth differentiation factor 15 (GDF15) [47–49], among others. The role of these hormones in promoting lipoatrophic mice hypermetabolism will be addressed in future studies.

In conclusion, lipoatrophy hypermetabolism at thermoneutral conditions is driven in part by an increase in thermic effect of food due to hyperphagia and basal metabolic rate due to enhanced liver *de novo* lipogenesis and 5-fold increase in liver mass. This enhanced energy expenditure widened the thermoneutral zone and caused hyperthermia in lipoatrophic mice at thermoneutral conditions. Further studies should address the contribution other thermogenic processes and hormones to lipoatrophic mice hypermetabolism.

## Acknowledgements

This work was supported by grants from Sao Paulo Research Foundation (FAPESP #15/19530-5, 19/01763-4, 20/04159-8, 22/11234-1, 25/04262-7) and Brazilian National Council for Scientific and Technological Development (CNPq, #303784/2022-9) to WTF. ÁSP, BFL, ÉC, TSV, JVF, ABP, NMP, EVMP, MAA-H, ACPB, LS, MM, LBG, ABC-F and TEO were recipients of fellowships from FAPESP (#22/02123-1, 21/14419-0, 26/01093-2, 20/16656-6, 23/04753-5, 24/01632-5, 24/09406-4, 23/04509-7, 24/12973-8, 24/13040-5, 23/17140-1, 24/11195-1, 24/16241-1, 25/14824-2, 21/10153-5, 23/12767-6). LPS-J, MLEM and SS were recipients of fellowship from Coordenação de Aperfeiçoamento de Pessoal de Nível Superior (CAPES) 88887.673950/2022-00, 8887.084152/2024-00, 88887.251830/2026-00. EMS and RJM were recipients of fellowship from CNPq 144006/2025-1 and 144006/2025-1.

## Notes

### Competing Interest Statement

The authors have declared no competing interest.

